# Haploid-phased chromosomal telomere to telomere genome assembly of *Uncaria rhynchophylla* accelerating gene mining on the biosynthesis of medicinal alkaloids

**DOI:** 10.1101/2024.05.30.596046

**Authors:** Tao Hu, Lei Duan, Liyang Shangguan, Qingshi Zhao, Ye Hang, Xue Li, Nngxian Yang, Fulin Yan, Qiuyu Lv, Liu Tang, Miao Liu, Wei Qiang, Xincun Wang, Xuewen Wang, Mingsheng Zhang

**Author notes:** Corresponding authors (M. Zhang); (X. Wang).

## Abstract

Recorded traditionary medicinal plants contain important resources, including many natural medicinal alkaloids, for new medicine discovery and development. *Uncaria* genus is such a woody plant, with a high medicinal value in alkaloids, e.g. (iso)rhynchophylline. Natural alkaloids’ contents usually vary between germplasms and are affected by the growth environment, which requires a genomics solution to understand the genetic and environmental factors that influence alkaloid production in more detail. Here, we have dissected the haploid-resolved chromosomal T2T genome assembly of *Uncaria rhynchophylla* with a size of ∼634 Mb and contig N50 of 26 Mb using PacBio HiFi long-reads and Hi-C and anchored the contigs on 22 pairs of confirmed chromosomes. This genome contains 56% repeat sequences and ∼2,9000 protein-encoding genes. *U. rhynchophylla* diverged from a common ancestor shared with *Coffea* around 20 million years ago and contains expanded and contracted gene families associated with secondary metabolites and plant-associated defenses. We constructed the pathway for (iso)rhynchophylline biosynthesis with genes mined from the genome and comparative transcriptomes. 53 alkaloids out of 2,578 metabolites were identified in (iso)rhynchophylline biosynthesis, where eight differentially expressed genes were the key for regulating the catalytic steps leading to alkaloid abundance difference between tissues. The chromosome-level genome and pathways of (iso)rhynchophylline constructed in this study provide a genetic basis and guidance for further breeding improvements and the development of pharmaceutical alkaloids.

## Introduction

The *Uncaria* genus is a diverse group of plants with significant medicinal value in alkaloids (Lopes et al., 2019). The *Uncaria* genus, part of the Rubiaceae family, is a flowering plant that consists of about 40 species (Castro et al., 2023). The distribution of these species is pantropical, with most species native to tropical Asia, three from Africa and the Mediterranean, and two from the neotropics (Wikipedia, 2024). A comparative and phylogenetic analysis of the complete chloroplast genomes of six *Uncaria* species showed that the genus *Uncaria* is a monophyletic group, belonging to the tribe Naucleeae (Dai et al., 2023). It also clarified that *U. sinensis* is not a variant of *U. rhynchophylla* and found that *U. rhynchophylloides* and *U. rhynchophylla* are not the same species, which suggests complexity (Dai et al., 2023). Around 12 species of *Uncaria* are distributed and are commonly known as “Gou Teng” in traditional Chinese medicine. *U. rhynchophylla* is the major species used for herb medicine.

The active medicinal compounds are reported as indole and oxindole alkaloids. Some alkaloids identified from *U. rhynchophylla* include rhynchophylline (RIN), isorhynchophylline (IRN), corynoxine, isocorynoxeine, hirsuteine, hirsutine, angustoline, angustidine and geissoschizine methyl ether. The estimated total alkaloid content in *U. rhynchophylla* is approximately 0.2% of the dry weight, among which RIN and IRN account for the major proportions (Zeng et al., 2021b). These alkaloids exert therapeutic effects on the cardiovascular and nervous system. For example, neuroprotective effects against neurodegenerative diseases such as Alzheimer’s disease through antioxidant, anti-inflammatory, and neuromodulatory activities (Kushida et al., 2021; Zeng et al., 2021b). Among those alkaloids including angustoline, angustidine, and isocorynoxeine were identified as the key compound regulating amyloid-β production, while corynoxine, isocorynoxeine, dihydrocorynatheine, IRN, and hirsutine were identified as key alkaloids that regulate tau phosphorylation (Zeng et al., 2021a; Zeng et al., 2021b).

The biosynthesis pathways of RIN in plants, particularly in *U. rhynchophylla*, are not fully understood, but some steps have been identified. The RIN biosynthesis pathway shares some early steps in other plants in the KEGG database, which includes the indole alkaloid biosynthesis pathway (map 00901), the monoterpenoid biosynthesis pathway (map 00902), and the tryptophan pathway (map 00400). After the condensation of tryptamine from the tryptophan pathway and secologanin from the monoterpenoid pathway, the strictosidine was synthesized, which was followed by further oxidation and modification (Yang et al., 2022). Stemmadenine and tabersonine are then the downstream compounds. In *U. rhynchophylla*, the RIN and IRN were derived from tabersonine, after further oxidization and methylation. Transcriptome analysis showed that ethylene and light intensity can regulate the RIN and IRN biosynthesis, in which assembled transcripts encoding G8O, IO, 7-DLGT, LAMT, TDC, and STR were mostly upregulated (Li et al., 2022; Wang et al., 2022).

To better understand the genomes and genes for the biosynthesis of RIN associated alkaloids, we used the most advanced high-accuracy HiFi long-sequencing reads to generate reference genome assembly, which improve genetic analysis (Aganezov et al., 2022). In this study, we report a de novo chromosomal haplotype-resolved genome assembly with HiFi and Hi-C data, annotation, biosynthesis gene mining, metabolomics analysis, and the gene expressional regulation on the accumulation of pharmacological alkaloids in *U. rhynchophylla.* Results generated from this study could accelerate genetic research like breeding improvement and medicinal compound mining for drug discovery.

## Results

### The chromosomal reference genome assembly

To explore the genome sequence and accelerate molecular gene-trait analysis, we generated a reference-quality haplotype-resolved chromosomal assembly for *U. rhynchophylla*. Karyotype analysis revealed that *U. rhynchophylla* is a diploid species with 22 pairs of chromosomes (2n = 2x = 44, **Figure 1**A). Genome size is estimated to be 600-650 Mb from k-mer analysis of 102.9 Gb (∼162x) 150-bp paired-end short reads from an Illumina platform. An initial haplotypic contig assembly has an N50 length of ∼26 Mb, which was assembled from 74 Gb (∼117x) PacBio HiFi long reads and phased with 81.5 Gb (∼128x) Hi-C short reads from the Illumina Novo-6000 platform (Supplementary Table S1). The assembled primary genome has a contig N50 length of ∼27 Mb (**Table 1**). After the Hi-C contact map analysis, over 95% of contig sequences in each phased assembly were mapped into 22 chromosomes, which matches the number of chromosomes identified in our karyotype analysis, suggesting a phased assembly (**Figure 1**B, C). The final two haplotypic assemblies have very close sizes of 630 Mb and 639 Mb (Supplementary Figure S1, Figure S2). Further comparison of the two haplotypic assemblies showed an identical sequence collinearity. Telomeres were presented on both ends of all 21 pseudo-chromosomes except one telomere on chromosome 9. Sixteen chromosomes have gapless telomere to telomere assembly (**Figure 1**D). The assembly has a score of 98.4% gene completeness and no missing genes against the orthologs database embryophyta_odb10 in Benchmarking Universal Single-Copy Orthologs (BUSCO) (**Table 1**).

**Figure 1.**
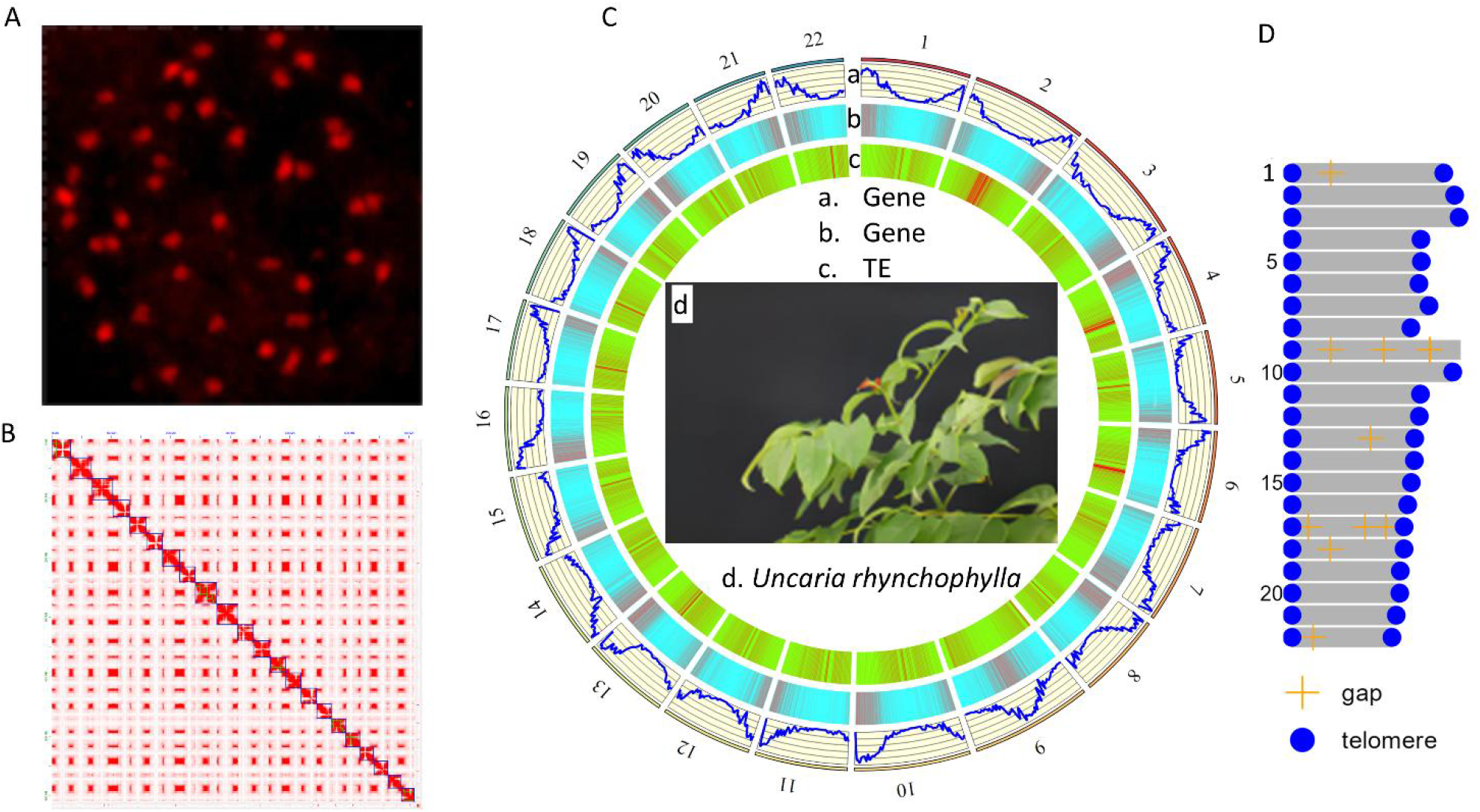
Chromosomes, hi-C map, and genome plots of assembly for U. rhynchophylla. **A.** The karyotype shows 22 pairs of chromosomes in *U. rhynchophylla*. **B.** Hi-C contact map showing the anchoring of contigs into 22 chromosomes in haplotype genome assembly. **C.** The circular plot shows the gene density and transposon element (TE) density in the genome assembly and plant image. **D**. The illustration shows the telomeres and gaps on 22 chromosomes.

**Table 1.** Statistical summary of U. rhynchophylla genome assembly.

|  | Item | N50 (bp) | L50 | N90 (bp) | L90 | Length (bp) |
| --- | --- | --- | --- | --- | --- | --- |
| Assembly | Hap1 contig | 26,050,781 | 11 | 11,603,464 | 24 | 639,240,331 |
| (contig v0.4) | Hap2 contig | 26,316,424 | 11 | 9,507,307 | 25 | 630,397,526 |
|  | Primary |  |  |  |  |  |
|  | contig | 27,618,998 | 11 | 23,097,658 | 21 | 650,186,565 |
| (chromosome |  |  |  |  |  |  |
| v1.0) | Hap1_chr | 26,354,137 | 11 | 23,049,914 | 21 | 639,241,131 |
|  | Hap2_chr | 27,379,396 | 11 | 23,181,917 | 21 | 630,398,626 |
|  |  |  |  |  | Mean | 634,819,879 |
|  | eukaryota_odb10 |  |  |  |  |  |
| BUSCO |  |  |  |  | Count | Percent |
|  | Complete BUSCOs |  |  |  | 251 | 98.40% |
|  | Complete and single-copy BUSCOs |  |  |  | 179 | 70.20% |
|  | Complete and duplicated BUSCOs |  |  |  | 72 | 28.20% |
|  | Fragmented BUSCOs |  |  |  | 4 | 1.60% |
|  | Missing BUSCOs |  |  |  | 0 | 0.00% |
|  | Total BUSCO groups searched |  |  |  | 255 | 100% |

### Repeat sequence and gene annotation

To investigate the repeat sequences in the genome, the combined methods of *de novo* and homology search were used. We identified that 355 Mb (56.34%) of the genome assembly belongs to repetitive sequences. Of those, Copia (7.99%) and Gypsy (18%) in the LTR element category are the most abundant ones (**Table 2**), which is normal in plant genomes with similar size. 29,049 protein-encoding genes were predicted in the assembly with *ab initio* prediction, homology search, and RNA-seq transcript evidence from the pooled root, stem, and leaf, etc. (Supplementary Table S2). 20,821 (72%) genes have functional annotation in the databases of GO, KEGG, PFAM, and COG (Supplementary Table S4). Besides, 8,062 small RNAs were annotated in the assembly.

**Table 2.** Annotation of repeat sequence in the U. rhynchophylla genome.

|  | Element count* | Length (bp) | Percent of genome |
| --- | --- | --- | --- |
| Retroelements | 202480 | 198948917 | **31.56%** |
| SINEs: | 265 | 381206 | 0.06% |
| Penelope: | 105 | 36695 | 0.01% |
| LINEs: | 12412 | 12012012 | 1.91% |
| R1/LOA/Jockey | 1019 | 769494 | 0.12% |
| RTE/Bov-B | 1298 | 521558 | 0.08% |
| L1/CIN4 | 10095 | 10720960 | 1.70% |
| LTR elements: | 189803 | 186555699 | **29.59%** |
| **Ty1/Copia** | 50651 | 50382590 | 7.99% |
| **Gypsy/DIRS1** | 99002 | 113475263 | 18.00% |
| Retroviral | 1199 | 147657 | 0.02% |
| DNA transposons | 18587 | 7982905 | 1.27% |
| hobo-Activator | 2914 | 1819121 | 0.29% |
| MULE-MuDR | 6561 | 2534544 | 0.40% |
| Tourist/Harbinger | 1194 | 602400 | 0.10% |
| Rolling-circles | 2086 | 2230577 | 0.35% |
| Unclassified: | 369570 | 148212956 | 23.51% |
| **Total interspersed repeats:** |  | **355181473** | **56.34%** |

### Evolutionary and comparative genomic analyses

To reveal the evolutionary phylogeny of *U. rhynchophylla*, a comparative analysis was conducted by using the protein-coding genes from five additional species including Arabidopsis (Kang et al., 2023), grape (*Vitis*) (Shi et al., 2023), two close taxonomy-related coffee species (Salojärvi et al., 2024), and rice (*Oryza*) (Stein et al., 2018) as the outgroup. We identified 31,125 orthologs and 1,944 paralogs in *U. rhynchophylla*. A phylogenetic tree was constructed from 1,860 conserved single-copy ortholog genes, where *U. rhynchophylla* was clustered into the same clade and diverged from a common recent ancestor of *Coffea* at ∼ 20 million years ago, slightly earlier than the divergency of two Coffea species (**Figure 2**A), as expected in NCBI taxonomy database. 1,269 gene families were expanded, and 1,765 gene families were contracted, which are smaller than those in *Vitis* (**Figure 2**A). The expanded genes were enriched in response to diterpene (GO:1904629), sesquiterpenoid catabolic process (GO:0016107), modulation by host of viral process (GO:0044788), modulation by host of symbiont process (GO:0051851), positive regulation of defense response to insect (GO:1900367). 11,197 common protein-encoding gene families were identified among the *A. thaliana*, *U. rhynchophylla*, *V. vinifera,* and *C. arabica*. Fewer unique genes in *U. rhynchophylla* were present than other species investigated here (**Figure 2**B). The 4DTv distribution of paralogs indicated a well-kept trace of ancient whole genome duplication events in the *Uncaria* (**Figure 2**C).

**Figure 2.**
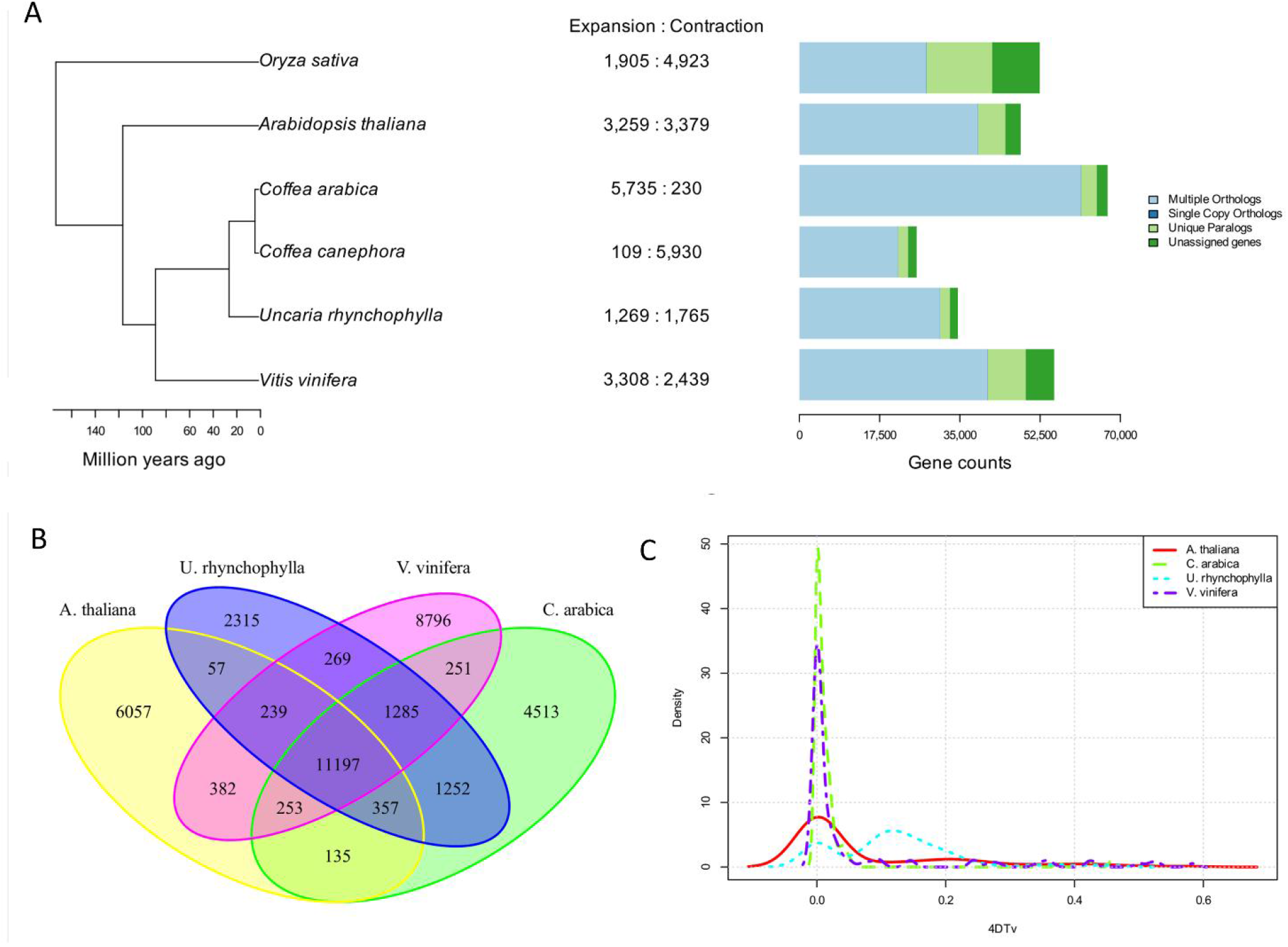
Comparison analysis of phylogenetic tree and gene family in U. rhynchophylla. **A.** Phylogenetic tree of *U. rhynchophylla, g*ene family expansions, and contractions, clusters of orthologous and paralogous gene families. **B.** Venn diagram representing the distribution of shared genes between *U. rhynchophylla* and other plants. **C.** Distribution of paralog-pair 4DTv density indicating whole-genome duplications.

### Gene expression and tissue-specific regulation in *U. rhynchophylla*

The expressed genes in the stem, leaf, and root were investigated with deep RNA-Seq paired-end reads in three independent experiments. In total, 90,194 transcripts from 32,206 genes, an average of 2.8 transcripts per gene, were found to be expressed among all tissues. Of those, 29,027 expressed genes were supported by at least two RNA-Seq reads. Differentially expressed genes (DEGs) were identified with the criteria of > 2 folds of change in transcript abundance and adjusted *p* value < 0.001 (**Figure 2**A, Supplementary Table S3). We compared the expression of DEGs with a focus on medicinal stems relative to other tissues (**Figure 2**B). Compared with those in leaves, 287 up-regulated and 227 down-regulated DEGs were identified in stems (Supplementary Figure S5). Similarly, relative to those in roots, 207 up-regulated and 296 down-regulated DEGs were discovered in stems (Supplementary Figure S4). More DEGs, 517 up-regulated and 656 down-regulated, were found in leaves compared with roots (**Figure 2**C, Supplementary Figure S6). The expression patterns of DEGs identified between stems and leaves also had very similar expressional patterns between stems and roots (**Figure 2**B). Interestingly, 104 DEGs exhibited differential expression between any two tissues (**Figure 2**C). The difference in DEG counts suggests more similar expression patterns between stem and leaves than those between leaves and roots. Further pathway and functional analysis revealed that some DEGs are involved in the RIN closely associated pathways. The DEG-associated pathways and annotations were summarized and linked to the KEGG database in supplementary data (Supplementary Table S6, S7, and S8). Among the DEGs from the stem and root comparison, one encodes cytochrome P450 [(KEGG id ko00199)] as an oxidoreductase in the precursor biosynthesis of terpenoids at upstream of RIN. Two DEGs were in the RIN-associated tryptophan pathway, encoding caffeic acid 3-O-methyltransferase / acetylserotonin O-methyltransferase [EC:2.1.1.68, 2.1.1.4, KEGG id K13066] and 2-oxoglutarate-dependent dioxygenase [EC:1.14.11.-, KEGG id K23947]. Further DEGs integrated KEGG pathway enrichment across tissues showed that the DEGs were enriched in the biosynthesis of secondary metabolites, metabolic pathways, and plant defenses (**Figure 2**D).

**Figure 3.**
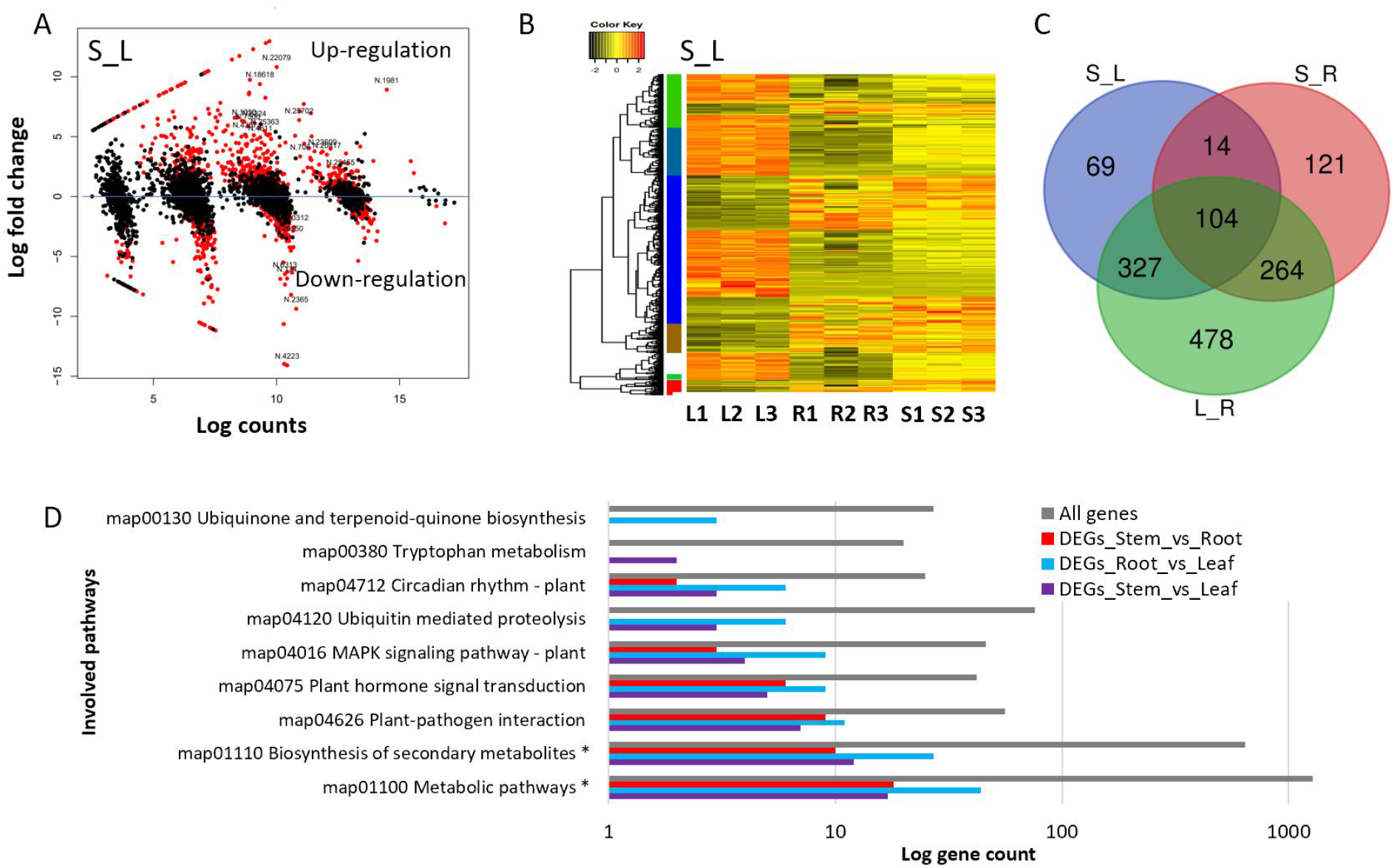
DEGs between tissues and biological pathways involved in U. rhynchophylla. **A.** A volcano plot showing the differentially up- and down-regulated gene expression between stem (S) and leaf (L). **B.** The expressional patterns of DEGs identified between stem and leaf. **C.** The Venn diagram shows the common and unique DEGs. **D.** A comparison of enriched pathways for all DEGs between tissues of stem, root (R), and leaf. * represents a statistical significance (*p* < 0.05) in a hypergeometric test.

### Secondary metabolites in tissues of *U. rhynchophylla*

To detect the natural products in *U. rhynchophylla*, metabolites in leaves, stems, and roots, the same tissues as those used in RNA-Seq analysis, were investigated with HPLC-MS. In total, 2,578 metabolites were identified, with 2,442 having a trackable identifier in the human metabolome database (HMDB) and 1,179 in the KEGG database. Principal component analysis revealed that the abundance patterns of compounds in the leaves are more closely correlated to those in stems than to those in roots, which is consistent with transcriptomic patterns. 51 detected metabolites belong to the alkaloid superclass of the HMDB, including those in the upstream and immediate steps of IRN biosynthesis in the indole alkaloid biosynthesis pathway, the monoterpenoid biosynthesis pathway, and the tryptophan pathway. We further described these compounds in late sections.

### Expressional regulation on biosynthesis pathway of medicinal alkaloid metabolites

To mine the genes responsible for the medicinal alkaloids in *U. rhynchophylla*, we reconstructed a biosynthesis pathway map by integrating the monoterpenoid biosynthesis (map 00902), the indole alkaloid biosynthesis (map 00901) of plants from the KEGG database (www.kegg.jp), and others from published literature (St-Pierre and De Luca, 1995; Li et al., 2022; Wang et al., 2022; Yang et al., 2022). The combined pathway starts from geranyl-diphosphate to produce loganin with a cyclopentane ring in the map 00902 (**Figure 4**). Then in map 00901, the loganin is cleaved in a further cytochrome P450-dependent step, leading to secologanin, which provides the carbon skeleton for many bioactive indole alkaloids. The secologanin combines tryptamine derived from the amino acid tryptophan to generate the strictosidine. Strictosidine is further used to generate other alkaloid intermediates and the tabersonine, which is believed to be further oxidized to produce RIN and IRN with a potential involvement of the cytochrome P450. The RIN and IRN were 5-10 times higher than other alkaloids in *U. rhynchophylla*. The highest RIN amounts were found in the leaf and stem. The detailed steps from tabersonine to RIN and IRN are still unknown (**Figure 4**). In addition, we found several previously unreported alkaloids in *U. rhynchophylla* in this pathway. Among them, polyneuridine aldehyde, gentiopicrin, and camphor had very low abundance in the root but high abundance in the leaf and stem (Supplementary Figure S3).

**Figure 4.**
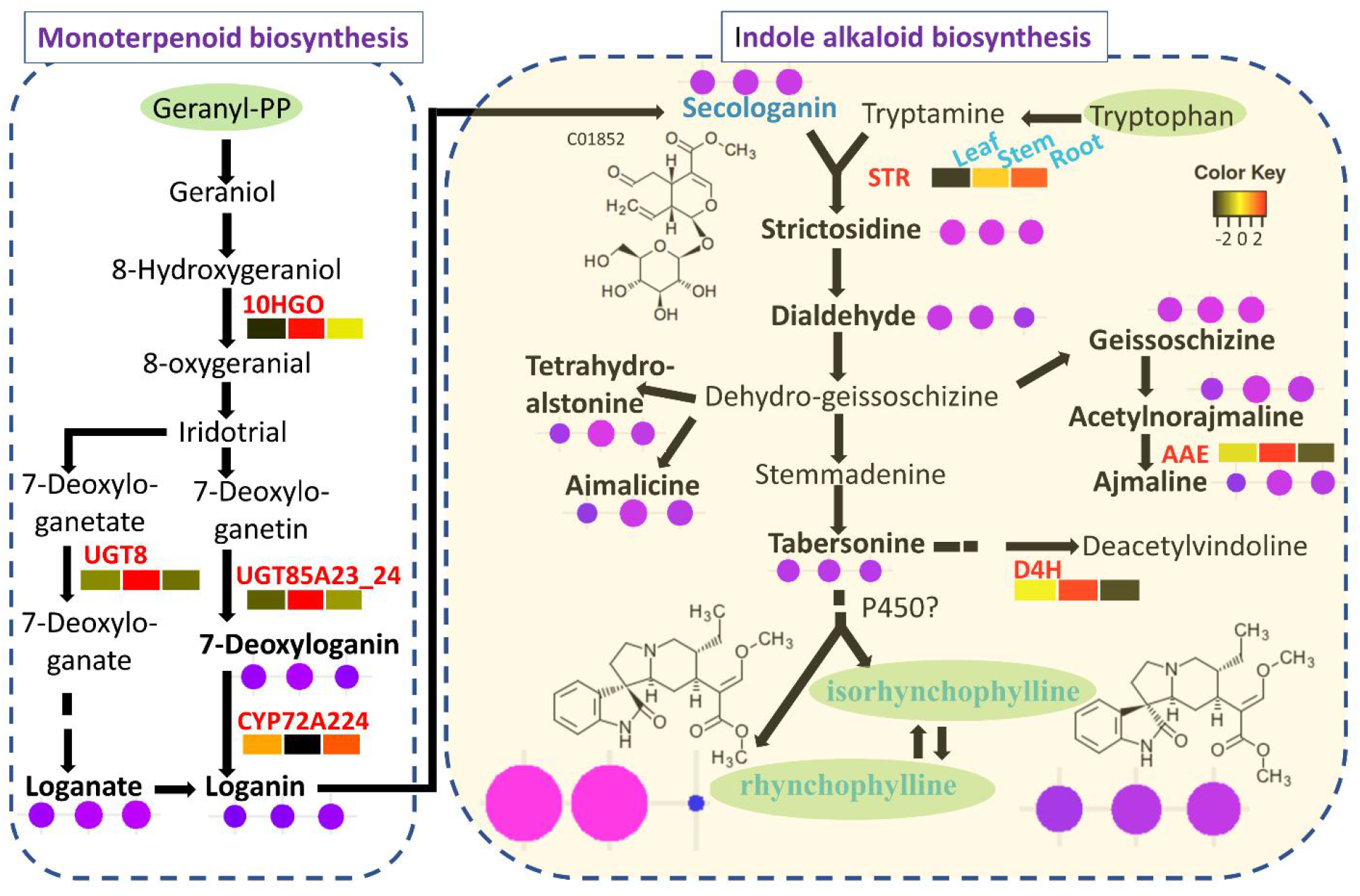
Biosynthesis pathway, gene expression, and alkaloids toward (iso)rhynchophylline.

The illustration shows the currently integrated knowledge of the biosynthesis of (iso)rhynchophylline. The differentially expressed genes (DEGs) described in **Table 3** in associated catalytic steps were highlighted in red next to the arrow. The expression levels in FPKM of DEGs in the leaf, stem, and root were shown in a heatmap in a rectangle shape. The levels were shown after the log2 transformation of the mean values from three experiments. The abundance of metabolite compounds in ug/g weight was plotted in a circle in proportion to the abundance levels next to each compound in tissue order of leaf, stem, and root. The abundance shown here is the mean of values from three experiments.

Most of the alkaloids in the integrated pathway were detected in our metabolome dataset (Supplementary Figure S3). Of those, the important alkaloids were highlighted in bold fonts (**Figure 4**), including secologanin, tabersonine, targeted (iso)rhynchophylline, etc. Genes for the most known steps were also identified in our transcriptomes or genes in the assembly. To reveal the association between the gene’s role and metabolites, we highlighted the DEGs in the pathway in red and presented their expression levels in a rectangle heatmap, along with the metabolite abundance in circles. Among eight identified DEGs in the integrated pathway (**Table 3**), three DEGs (*10HGO* also termed cytochrome P450, *UGT8,* and *UGT85A23-24*) exhibited a higher expression in the stem than in other tissues during steps of the monoterpenoid biosynthesis, while two DEGs catalyzing the steps from loganin to strictosidine during the transition from monoterpenoids and indole alkaloids expressed much higher in root than other tissues (**Figure 4**). Four DEGs in the indole alkaloid biosynthesis were identified and three of them were involved in (iso)rhynchophylline biosynthesis (**Table 3**). Expression of AAE, D4H, and NMDH was much higher in the stem than in other tissues. A positive correlation was found between loganin and gene *CYP72A224,* ajmaline, and gene *AAE.* No correlation was found between strictosidine and gene *STR*. For other metabolites, the associated gene expression did not exhibit differential expression (Supplementary Figure S3). These results suggest that differential expression in the catalytic enzyme encoding genes should have different roles in regulating alkaloid biosynthesis in a tissue-specific way. These also suggested that early steps mostly happened in the stem transition steps happened in the root and later steps are in the stem again.

**Table 3.** DEGs in the (iso)rhynchophylline biosynthesis.

| Map | KEGG ID | Symbol | Enzyme name | Enzyme ID |
| --- | --- | --- | --- | --- |
| 00902 | K15095 | NMDH | (+)-neomenthol dehydrogenase<sup>#</sup> | [EC:1.1.1.208] |
| 00902 | K23276 | CYP72A224 | Cytochrome P450, 7-deoxyloganate 7-hydroxylase | [EC:1.14.14.85] |
| 00902 | K23232 | 10HGO | Cytochrome P450, 8-hydroxygeraniol dehydrogenase | [EC:1.1.1.324] |
| 00902 | K21374 | UGT85A23_24 | 7-deoxyloganetin glucosyltransferase | [EC:2.4.1.324] |
| 00902 | K21373 | UGT8 | 7-deoxyloganetic acid glucosyltransferase | [EC:2.4.1.323] |
| 00901 | K21026 | AAE | acetylajmaline esterase | [EC:3.1.1.80] |
| 00901 | K12697 | D4H | deacetoxyvindoline 4-hydroxylase | [EC:1.14.11.20] |
| 00901 | K01757 | STR | strictosidine synthase | [EC:3.5.99.13] |

\* Not shown on the map because it is in a far branch.

## Methods

### Plant materials and karyotype analysis

*Uncaria rhynchophylla* has been used in this study. Fresh root tips from the same plant *U. rhynchophylla* for genome sequencing were used to observe the chromosomes. The root tips were treated and fixed in a solution of acetic acid and alcohol to preserve the cells and chromosomes.

Then, the root tip was squashed and stained with aceto-orcein followed by observation under a microscope. The number of chromosomes was counted for more than ten cells to determine the final number of chromosomes.

### Genomic DNA extraction, sequencing, and genome assembly

High molecular weight genomic DNA was extracted from the leaves of one *U. rhynchophylla* plant with the previously described method (Devos et al., 2023). For the genome survey, the genomic DNA was used to construct a sequencing library and sequenced on an Illumina NovaSeq 6000 platform using a whole genome shotgun sequencing strategy and standard sequencing protocols. The generated 150-bp paired-end reads, ∼102 G bp, were then used for subsequent genome size estimation.

The PacBio HiFi long-read sequencing strategy was used to generate sequences for genome assembly. The same batch of genomic DNA was used to construct the SMRTbell l5-kb library with the PacBio SMRTbell Express Template Prep Kit 3.0 according to manufacturer recommendations and then sequenced on a PacBio Sequel IIe platform with the Sequel II Sequencing Kit 2.0 and Sequel Binding Kit 3.2. Two SMRT cell runs were conducted. After sequencing, the resulting HiFi reads were generated with SMRT Link (version 11.0) software with default settings. In total, ∼74 G bp HiFi reads were generated with an average length of ∼17 kb.

### Hi-C sequencing data generation

The leaf tissue from the plant for genome sequencing was used. The Chromatin Conformation Capture (Hi-C) library was constructed from enzyme DpnII (NEB, USA) digestion of cross-linked DNA and sequenced to produce 150 bp paired-end reads with an Illumina Novaseq 6000 machine following the manufacturer’s sequencing protocols. In total, ∼82 G bp clean data were obtained.

### Genome assembly and chromosome anchoring

The HiFi long-read data were filtered to get high-quality sequences, which have a minimum length of five kb and average read quality higher than Q30 with our read processing tools (https://github.com/XuewenWangUGA/SeqReadsProcessing). The filtered sequences were used to build the contigs and then further scaffold with Hi-C data with Hifiasm (version 0.17). The peak Kmer coverage and noise cut-off values were manually curated to warranty assembly parameters and used the 51 mer peak distribution information during assembling. Two haploid assemblies were obtained, each for one parental inherited haplotypic genome set. Hi-C reads were preprocessed with Trimomatic (version 0.33) and then mapped to the contigs with BWA (version 0.7.17) as described previously in detail (Wang et al., 2019). YaHS (version 1.0)(Zhou et al., 2022) and Juicer tools (the version built in YaHS) were used to build the first version of anchored scaffolds and contigs, followed by purging duplicated contigs out of unanchored contigs. Juicebox (version 1.11) was used to validate and correct the chromosome anchoring and generate the final assembly with the Juice assembly tool (version 1.89)(Durand et al., 2016). The number of chromosomal scaffolds was compared with that in the karyotype result. Hi-C anchoring was conducted for both haploids and achieved scaffolds were well matched each other and had the same orientation in the DNA sequence alignment. The chromosome IDs (e.g., chr1) were ordered by one-phased haplo-assembly first based on the Hi-C heat map from left top to right bottom followed by the Pan-Sol specification (https://github.com/pan-sol/pan-sol-spec/blob/main/v0.md). Both long and short sequencing reads were mapped back to the assembly to ensure good coverage on all assembled sequences. The gene completeness in the assembly was evaluated with BUSCO orthologs in the specified database eukaryote_odb10. The telomeres were identified with quartet (Lin et al., 2023) and motif search for TTAGGG/CCCTAA at the end of each assembly.

### Repeat sequence annotation and gene annotation

The repeat sequence was annotated with structure- and homology-based methods slightly modified from that in our previous report (VanBuren et al., 2020). Briefly, first, the SSR in the genome sequence was masked with GMATA (version 2.2)(Wang and Wang, 2016). Then masked genome was used to mine *de novo* LTRs with Genome tools, LTR-finder, and (version 2.9.0) (Ou and Jiang, 2018). The Dfam (version 3.7, https://www.dfam.org/ (Storer et al., 2021) TE models were used for homology search with RepeatModeler 2 (version 2.0.4)(Flynn et al., 2020). Repeat sequences from both methods were classified via CD-HIT (version 4.8.1) to generate a customized library for RepeatMasker (version 4.1.5) to analyze interspersed TEs in the whole genome assembly (Smit, 2015).

Gene annotation was conducted by combining *ab initio* prediction, RNA-Seq evidence, and homolog search with Braker pipeline (version 3) integrated with AUGUSTUS, gmst, and GeneMark.hmm3 (Hoff et al., 2019; Gabriel et al., 2024).

### RNA extraction and RNA-Seq transcriptome analysis

For genome annotation, the one RNA-Seq dataset was generated from a pool of fresh tissues of leaves, stems, hooks, buds, roots, etc. from the plant used for genome sequencing. For tissue-specific expression comparisons, RNA-Seq data were produced from RNA extracted from different tissues. In detail, RNA was extracted separately from each of the three types of fresh tissues of the leaf, root, and stem. Three plant individuals around three years old were used in one pool in each experiment. The leaves and stems grown from the current year were sampled. Three independent experiments were conducted. Transcriptome data were generated from the TrueSeq mRNA library on an Illumina NovaSeq 6000 platform following manufacturer recommendations. In total, nine RNA-Seq datasets were obtained for three types of tissues. The reads are in 150-bp paired-end format. Raw reads were trimmed, filtered, and quantified with the genome assembly as the reference as described in detail in our previous publication (Wang et al., 2019). DEGs between tissues were identified with DESeq2 with at least 2 folds of change in abundance and adjusted P value less than 0.001 following previous detailed methods (Wang et al., 2019). Three independent experiments were conducted and sequenced separately. In total, nine RNA-Seq datasets were generated, each with 7-9 G bp reads. Assembled transcripts were supported by at least 2 reads that had a length longer than 200 bp and were kept for subsequent analysis. Gene expression levels were presented in fragment per kilobase per million reads (FPKM). Gene ontology (GOs) and pathways of transcripts were annotated against databases (http://eggnog5.embl.de/#/app/home) with EggNOG mapper (v2.1.12), KEGG database (accessed on 2023 Sept), and Panther version 17 (https://pantherdb.org/).

### Phylogenetic and comparative analysis

To analyze the phylogenetic relatedness, an investigation of taxonomy was performed at NCBI (https://www.ncbi.nlm.nih.gov/taxonomy), and a literature search was conducted to determine closely related species with high-quality genome assembly data. Selected genome assembly datasets were downloaded. These genomes include *Arabidopsis thaliana* (v10), *Vitis vinifera* (grape, v2.1), *Coffea arabica* (v0.5), *Coffea canephora* (v0.5), and *Oryza sativa* (rice, v7.0) as the outgroup from Phytozome (https://phytozome-next.jgi.doe.gov/). Orthologs of protein-coding genes were aligned with diamond (v2.1.9, options: --more-sensitive -p 1 --quiet -e 0.001 --compress 1) and identified with OrthoFinder2 v 2.5.5 (Emms and Kelly, 2019). The phylogenetic tree was constructed with all single-copy orthologs and with calibration of the estimated time from Timetree of Life (https://timetree.org). Orthologs and paralogs were annotated and classified again by PANTHER HMM gene family models (v17.0) to obtain gene ontology (GO) information and family classification information (Mi et al., 2019). Gene expansion and contraction were analyzed with CAFÉ (v5, options: -p 0.05 -t 8 -r 10,000 -filter) (De Bie et al., 2006). GO enrichment analysis was conducted for expanded and contracted gene families with the hypergeometric test (*P* < 0.05). 4DTv distribution was analyzed from OrthoFinder2 identified paralogs to indicate whole genome duplication events.

### Metabolite detection and analysis

Untargeted metabolites were profiled for leaves, stems, and roots, which were also used for RNA-Seq transcriptome analysis. Each tissue was collected from three biological plants. Three independent experiments were conducted to detect untargeted metabolites. Fresh tissues were frozen immediately in liquid nitrogen before extraction. For each sample, 50 mg powder was extracted with 1.2 ml 80% methanol solution (volume methanol : H2O = 4 : 1) overnight at 4 °C. The extracts were filtered through a membrane (SCAA-104, 0.22-mm pore size) after centrifugation at 12,000 rpm for 10 min. Metabolite analysis for 20 μl supernatants was performed by ultraperformance liquid chromatography and tandem mass-mass spectrometry on the UHPLC-Q Exactive HF-X (Thermo Fisher Scientific, USA). Chromatographic conditions were set as follows. The chromatographic column is ACQUITY UPLC HSS T3 (100 mm × 2.1 mm i.d., 1.8 µm; Waters, Milford, USA); Mobile phase A is 95% water + 5% acetonitrile (containing 0.1% formic acid); Mobile phase B is 47.5% acetonitrile + 47.5% isopropanol + 5% water (containing 0.1% formic acid); The injection volume is 3 μL, and the column temperature is 40 °C. The data and compounds were retrieved with Progenesis QI (Waters Corporation, Milford, USA) from the elution profile against HMDB (http://www.hmdb.ca/, https://metlin.scripps.edu, accessed on 2023 Sept) and KEGG database. RIN and IRN were measured with Agilent 1100 HPLC Compass C18 (2) 250 mm*4.6 mm, 5 μm by using a standard curve built from the dilution series of the purified RIN and IRN compound purchased from Shanghai Yuanye.

### Software parameters

The parameters in all mentioned analyses were set to the default settings except those claimed specifically in our methods.

## Discussion

### Constructing chromosomal-level genome assembly of medicinal plants

Available genome assembly is the basis for further genetic analysis and pharmaceutical application of natural products. More available genomes are expected for refined germplasm and genetic manipulation for medicinal plants. In the study, we resolved the chromosome number 2n = 2x = 44 in *U. rhynchophylla* and generated chromosomal genome assembly with 22 pseudochromosomes in each phased haplotype. The phased chromosomal assemblies of *U. rhynchophylla* fill in the blank of lacking genome assembly in *Uncaria* for the first time. Increasing herb medicinal plant genome projects, such as the *Stephania* genome (Leng et al., 2024), have emerged in the very recent two years with the new advances of NGS long-read sequencing technologies. Currently, several available herb medicinal plant genomes are reported with the sequenced long reads from the ONT platform, which is cheaper but with a high noise and low accuracy in bases leading to an error-prone in the genome assembly (Cheng et al., 2024). Here, the *U. rhynchophylla* has been assembled from PacBio HiFi long reads with the highest base quality >99.9% among all available long-read sequencing technologies (Wenger et al., 2019), indicating a better genome sequence quality. The N50 contig of 26 Mb in length is enough to fully span every single gene and transposon element. The genome assembly we reported here is of high quality, demonstrated in our highly consistent parental phased haplotype 1 and haplotype 2 assemblies with an expected 22 chromosomes and higher gene completeness in BUSCO score based on lineage-specific orthologs (Manni et al., 2021). The haplotype-resolved assembly paves a new way to understand heterozygosity in two sets of haplotypic genomes and the roles of allelic variation in genes or proteins. Therefore, our results of the *U. rhynchophylla* genome advance the genetic and molecular analysis of medicinal plants.

### Enriched resources of gene and alkaloid biosynthesis

Natural products from herb plants have drawn great attention in new pharmaceutical development and application. Alkaloids are one of the largest categories of such natural products (Atanasov et al., 2021; Wang et al., 2021; Zeng et al., 2021b). Here we detected 51 alkaloids in addition to (iso)rhynchophylline in *U. rhynchophylla*, suggesting a high diversity of alkaloids. These detected compounds provide a landscape of secondary metabolites for further pharmaceutical development. Secondary metabolites are catalyzed by enzymes, which are encoded by genes. The genes annotated in the genome assembly, transcriptome, and metabolite pathways enrich the resources for gene regulation and functional research for the secondary metabolites for the community. Here the constructed pathway for (iso)rhynchophylline elucidates the genes and metabolites in the same network, which widens previous knowledge (Yang et al., 2022). The DEGs in the pathway will be the most important regulations responsible for the difference of metabolites between tissues, which may guide genetic control and medicinal material selection. Of course, the function of these DEGs may need further validation in vitro and in vivo experiments. The genes for the final step of (iso)rhynchophylline biosynthesis are still unknown but definitely in our expressed gene list. It may be the cytochrome P450 that catalyzed the oxidation and reduction reaction (Minerdi et al., 2023), but P450 is already involved in biosynthesis in the early steps. Thus, the P450 genes must have very diverged functions during evolution. The tissue-specific expressional patterns should play an important role in alkaloid accumulation, plant development, and adaptation although the molecular functions of these genes are not yet completely validated. In addition, expanded and contracted genes in secondary metabolites and plant defense responses will be good candidates for future investigation of the speciation of *U. rhynchophylla*. Existing knowledge suggests alkaloids are related to defenses (Atanasov et al., 2021; Wang et al., 2021; Zeng et al., 2021b). Combining the results of metabolites, genes, and expression, we hypothesize that the abundance of alkaloids in *U. rhynchophylla* may result from the diversity of genes responsible for the selection and adaptation to defenses.

### Genetic breeding improvement and medicinal tissue selection

The genetic study in *Uncaria* is very limited at the moment although it has been explored (Dai et al., 2023). The main reason is lacking genomic sequence reference. The genome assembly released here will facilitate the genetic diversity analysis. Several *Uncarcia* species are commercially grown on farms for medicinal raw materials. The molecular markers, e.g. simple sequence repeat (SSR) marker (Wang and Wang, 2016), can be developed using our genome sequence and genotyped in NGS data with tools like TRcaller (Wang et al., 2023) in many *Uncaria* species and current cultivars for breeding improvement. Given that the alkaloids accumulate differently in tissues, the results of high RIN in the leaf and stem of *U. rhynchophylla* provide clues for the preferred selection of these tissues for medicinal compound extraction. It is worth investigating whether this is the same in the other 40 species of *Uncaris* in the future.

## Data availability

All sequencing read data including PacBio HiFi long-read sequencing data, genome survey short-read reads, transcriptomic short-reads, and Hi-C reads are deposited at the National Center for Biotechnology Information (NCBI) short-read archives under accession number PRJNA1110988. The genome assembly and annotation files were stored in the GWH database at the National Genomics Data Center (NGDC) under accession number GWHESTA00000000.1 (https://ngdc.cncb.ac.cn/gwh).

## Author contributions

TH and XW designed the project; TH, LD, LYSG, XL, NY, FY, QL, LT and XW conducted experiments and data analysis; TH, QZ and YH collected the materials; XW guided the bioinformatics analysis. TH drafted the manuscript. MSZ and XW revised the manuscript. MSZ, ML, WQ supervised the project. All authors have read and approved the manuscript for publication.

