## Additional files for "Haploid-phased chromosomal telomere to telomere genome assembly of *Uncaria rhynchophylla* accelerating gene mining on the biosynthesis of medicinal alkaloids"

#### Contents

**Haploid-phased chromosomal genome assembly of *Uncaria rhynchophylla* accelerates gene mining on the biosynthesis of medicinal alkaloids** ....................................... **错误!未定义书签。**

**1. Supplementary Tables** .................................................................................................. 2

**2. Supplementary Figures** ................................................................................................ 11

### S. tables

### 1. Supplementary Tables

##########################################################################

**Stable 1. Summary of genome sequencing data**

| Read type | Length (G bp) | Estimated coverage |
|---|---|---|
| PacBio HiFi (15kb lib) | 74.1 | 117 |
| Hi-C (PE150) | 81.5 | 128 |
| Survey (PE150) | 102.9 | 162 |

**Stable 2. Summary of RNA-Seq transcriptome reads**

| Samples | #FASTQNAME | Clean Reads | Clean Bases | Format |
|---|---|---|---|---|
| Root1 | G-gen1_L2_388X88 | 22,571,030 | 6,771,309,000 | 150 paired-end |
| Root2 | G-gen2_L2_389X89 | 24,745,824 | 7,423,747,200 | 150 paired-end |
| Root3 | G-gen3_L2_390X90 | 21,126,444 | 6,337,933,200 | 150 paired-end |
| Stem1 | G-jing1_L2_391X91 | 25,271,235 | 7,581,370,500 | 150 paired-end |
| Stem2 | G-jing2_L2_392X92 | 24,282,488 | 7,284,746,400 | 150 paired-end |
| Stem3 | G-jing3_L2_393X93 | 28,197,440 | 8,459,232,000 | 150 paired-end |
| Leaf1 | G-ye1_L2_394X94 | 22,718,272 | 6,815,481,600 | 150 paired-end |
| Leaf2 | G-ye2_L2_395X95 | 24,109,269 | 7,232,780,700 | 150 paired-end |
| Leaf3 | G-ye3_L2_396X96 | 25,217,127 | 7,565,138,100 | 150 paired-end |

**Stable 3. Differentially expressed genes (>= 2 folds, P<0.001) between tissues**

| Comparison | Sample-control | Up-regulated genes | Down_regulated genes | Total DEGs |
|---|---|---|---|---|
| AA-BB | Stem v.s. Leaves | 287 | 227 | 514 |
| AA-CC | Stem v.s. Root | 207 | 296 | 503 |
| BB-CC | Leaves v.s. Root | 517 | 656 | 1173 |

**Stable 4. Summary of gene functional annotation**

| Annotation type | Count | Percent with annotation |
|---|---|---|
| eggNOG_Ogs/COGs | 20821 | 72% |
| GOs | 10039 | 35% |
| Kos in KEGG | 10409 | 36% |
| PFAMs | 19176 | 66% |
| Total genes | 29049 | 100% |

**Stable 5. Summary of tissue specific DEGs enriched pathways**

| Pathways | DEGs_Stem_vs_Leaf | DEGs_Stem_vs_Root | DEGs_Root_vs_Leaf | All genes |
|---|---|---|---|---|
| map01100 Metabolic pathways * | 17 | 18 | 44 | 1274 |
| map01110 Biosynthesis of secondary metabolites * | 12 | 10 | 27 | 641 |
| map04626 Plant-pathogen interaction | 7 | 9 | 11 | 56 |
| map04075 Plant hormone signal transduction | 5 | 6 | 9 | 42 |
| map04016 MAPK signaling pathway - plant | 4 | 3 | 9 | 46 |
| map04120 Ubiquitin mediated proteolysis | 3 | 0 | 6 | 76 |
| map04712 Circadian rhythm - plant | 3 | 2 | 6 | 25 |
| map00380 Tryptophan metabolism | 2 | 0 | 1 | 20 |
| map00130 Ubiquinone and terpenoid-quinone biosynthesis | 1 | 1 | 3 | 27 |

**Stable 6. Pathways and links of DEGs identified between stem and leaf**

| Listed links of stem_vs_leaf DEGs involved pathways |
|---|
| map01100 Metabolic pathways (17) |
| ko:K00876 udk; uridine kinase [EC:2.7.1.48] |
| ko:K00901 dgkA; diacylglycerol kinase (ATP) [EC:2.7.1.107] |
| ko:K01061 E3.1.1.45; carboxymethylenebutenolidase [EC:3.1.1.45] |
| ko:K01100 E3.1.3.37; sedoheptulose-bisphosphatase [EC:3.1.3.37] |
| ko:K01771 plc; 1-phosphatidylinositol phosphodiesterase [EC:4.6.1.13] |
| ko:K02641 petH; ferredoxin--NADP+ reductase [EC:1.18.1.2] |
| ko:K04123 KAO; ent-kaurenoic acid monooxygenase [EC:1.14.14.107] |
| ko:K06048 gshA; glutamate---cysteine ligase / carboxylate-amine ligase [EC:6.3.2.2 6.3.-.-] |
| ko:K12309 GLB1; beta-galactosidase [EC:3.2.1.23] |
| ko:K13066 COMT; caffeic acid 3-O-methyltransferase / acetylserotonin O-methyltransferase [EC:2.1.1.68 2.1.1.4] |
| ko:K13311 desVI; dTDP-3-amino-3,4,6-trideoxy-alpha-D-glucopyranose N,N-dimethyltransferase [EC:2.1.1.234] |
| ko:K16266 UGT85K; cyanohydrin UDP-glucosyltransferase [EC:2.4.1.-] |
| ko:K16839 hpxO; FAD-dependent urate hydroxylase [EC:1.14.13.113] |
| ko:K17872 NDC1; demethylphylloquinone reductase [EC:1.6.5.12] |
| ko:K19964 HIUH; hydroxyisourate hydrolase [EC:3.5.2.17] |
| ko:K21354 UGT94E5; beta-D-glucosyl crocetin beta-1,6-glucosyltransferase [EC:2.4.1.330] |
| ko:K22846 CS26; S-sulfo-L-cysteine synthase (O-acetyl-L-serine-dependent) [EC:2.5.1.144] |

map01110 Biosynthesis of secondary metabolites (12)
ko:K00901 dgkA; diacylglycerol kinase (ATP) [EC:2.7.1.107]
ko:K04123 KAO; ent-kaurenoic acid monooxygenase [EC:1.14.14.107]
ko:K12355 REF1; coniferyl-aldehyde dehydrogenase [EC:1.2.1.68]
ko:K13066 COMT; caffeic acid 3-O-methyltransferase / acetylserotonin O-methyltransferase [EC:2.1.1.68 2.1.1.4]
ko:K13071 PAO; pheophorbide a oxygenase [EC:1.14.15.17]
ko:K13311 desVI; dTDP-3-amino-3,4,6-trideoxy-alpha-D-glucopyranose N,N-dimethyltransferase [EC:2.1.1.234]
ko:K13492 ZOG1; zeatin O-glucosyltransferase [EC:2.4.1.203]
ko:K15095 E1.1.1.208; (+)-neomenthol dehydrogenase [EC:1.1.1.208]
ko:K15404 CER1; aldehyde decarbonylase [EC:4.1.99.5]
ko:K16266 UGT85K; cyanohydrin UDP-glucosyltransferase [EC:2.4.1.-]
ko:K17872 NDC1; demethylphylloquinone reductase [EC:1.6.5.12]
ko:K21354 UGT94E5; beta-D-glucosyl crocetin beta-1,6-glucosyltransferase [EC:2.4.1.330]

map04626 Plant-pathogen interaction (7)
ko:K13416 BAK1; brassinosteroid insensitive 1-associated receptor kinase 1 [EC:2.7.10.1 2.7.11.1]
ko:K13420 FLS2; LRR receptor-like serine/threonine-protein kinase FLS2 [EC:2.7.11.1]
ko:K13429 CERK1; chitin elicitor receptor kinase 1
ko:K13430 PBS1; serine/threonine-protein kinase PBS1 [EC:2.7.11.1]
ko:K13448 CML; calcium-binding protein CML
ko:K13453 PRF; disease resistance protein
ko:K13459 RPS2; disease resistance protein RPS2

map04075 Plant hormone signal transduction (5)
ko:K13416 BAK1; brassinosteroid insensitive 1-associated receptor kinase 1 [EC:2.7.10.1 2.7.11.1]
ko:K13422 MYC2; transcription factor MYC2
ko:K13464 JAZ; jasmonate ZIM domain-containing protein
ko:K14504 TCH4; xyloglucan:xyloglucosyl transferase TCH4 [EC:2.4.1.207]
ko:K14510 CTR1; serine/threonine-protein kinase CTR1 [EC:2.7.11.1]

map04016 MAPK signaling pathway - plant (4)
ko:K13416 BAK1; brassinosteroid insensitive 1-associated receptor kinase 1 [EC:2.7.10.1 2.7.11.1]
ko:K13420 FLS2; LRR receptor-like serine/threonine-protein kinase FLS2 [EC:2.7.11.1]
ko:K13422 MYC2; transcription factor MYC2
ko:K14510 CTR1; serine/threonine-protein kinase CTR1 [EC:2.7.11.1]

map01120 Microbial metabolism in diverse environments (4)
ko:K01061 E3.1.1.45; carboxymethylenebutenolidase [EC:3.1.1.45]
ko:K01100 E3.1.3.37; sedoheptulose-bisphosphatase [EC:3.1.3.37]
ko:K16839 hpxO; FAD-dependent urate hydroxylase [EC:1.14.13.113]
ko:K19964 HIUH; hydroxyisourate hydrolase [EC:3.5.2.17]

map04814 Motor proteins (4)
ko:K09290 TPM3; tropomyosin 3
ko:K10356 MYO1; myosin I
ko:K10357 MYO5; myosin V
ko:K10395 KIF4; kinesin family member 4

map04120 Ubiquitin mediated proteolysis (3)
ko:K03350 APC3; anaphase-promoting complex subunit 3
ko:K10594 HERC1; E3 ubiquitin-protein ligase HERC1 [EC:2.3.2.26]
ko:K10699 UBE1L2; ubiquitin-activating enzyme E1-like protein 2 [EC:6.2.1.45]

map04712 Circadian rhythm - plant (3)
ko:K12135 CO; zinc finger protein CONSTANS
ko:K16221 TCP21; transcription factor TCP21 (protein CCA1 HIKING EXPEDITION)
ko:K16222 CDF1; Dof zinc finger protein DOF5.5

map00380 Tryptophan metabolism (2)
ko:K13066 COMT; caffeic acid 3-O-methyltransferase / acetylserotonin O-methyltransferase [EC:2.1.1.68 2.1.1.4]
ko:K23947 DAO; 2-oxoglutarate-dependent dioxygenase [EC:1.14.11.-]

map00130 Ubiquinone and other terpenoid-quinone biosynthesis (1)
ko:K17872 NDC1; demethylphylloquinone reductase [EC:1.6.5.12]

**Stable 7. Pathways and links of DEGs identified between root and leaf**

| Listed link of Root_vs_leaf DEGs involved pathways |
|---|
| map01100 Metabolic pathways (44)<br>ko:K00079 CBR1; carbonyl reductase 1 [EC:1.1.1.184 1.1.1.189 1.1.1.197]<br>ko:K00310 SE; pyrimidodiazepine synthase [EC:1.5.4.1]<br>ko:K00333 nuoD; NADH-quinone oxidoreductase subunit D [EC:7.1.1.2] |

ko:K00472 P4HA; prolyl 4-hydroxylase [EC:1.14.11.2]
ko:K00487 CYP73A; trans-cinnamate 4-monooxygenase [EC:1.14.14.91]
ko:K00508 E1.14.19.3; linoleoyl-CoA desaturase [EC:1.14.19.3]
ko:K00827 AGXT2; alanine-glyoxylate transaminase / (R)-3-amino-2-methylpropionate-pyruvate transaminase [EC:2.6.1.44 2.6.1.40]
ko:K00876 udk; uridine kinase [EC:2.7.1.48]
ko:K00901 dgkA; diacylglycerol kinase (ATP) [EC:2.7.1.107]
ko:K00939 adk; adenylate kinase [EC:2.7.4.3]
ko:K01061 E3.1.1.45; carboxymethylenebutenolidase [EC:3.1.1.45]
ko:K01092 E3.1.3.25; myo-inositol-1(or 4)-monophosphatase [EC:3.1.3.25]
ko:K01100 E3.1.3.37; sedoheptulose-bisphosphatase [EC:3.1.3.37]
ko:K01470 E3.5.2.10; creatinine amidohydrolase [EC:3.5.2.10]
ko:K01771 plc; 1-phosphatidylinositol phosphodiesterase [EC:4.6.1.13]
ko:K02641 petH; ferredoxin--NADP+ reductase [EC:1.18.1.2]
ko:K04123 KAO; ent-kaurenoic acid monooxygenase [EC:1.14.14.107]
ko:K05714 mhpC; 2-hydroxy-6-oxonona-2,4-dienedioate hydrolase [EC:3.7.1.14]
ko:K06048 gshA; glutamate---cysteine ligase / carboxylate-amine ligase [EC:6.3.2.2 6.3.-.-]
ko:K06131 clsA_B; cardiolipin synthase A/B [EC:2.7.8.-]
ko:K06134 COQ7; 3-demethoxyubiquinol 3-hydroxylase [EC:1.14.99.60]
ko:K06287 yhdE; nucleoside triphosphate pyrophosphatase [EC:3.6.1.-]
ko:K06617 E2.4.1.82; raffinose synthase [EC:2.4.1.82]
ko:K06900 capV; cGAMP-activated phospholipase [EC:3.1.1.32 3.1.1.-]
ko:K08249 E4.1.2.11; hydroxymandelonitrile lyase [EC:4.1.2.11]
ko:K08729 PTDSS1; phosphatidylserine synthase 1 [EC:2.7.8.-]
ko:K10157 B3GAT2; galactosylgalactosylxylosylprotein 3-beta-glucuronosyltransferase 2 [EC:2.4.1.135]
ko:K12930 BZ1; anthocyanidin 3-O-glucosyltransferase [EC:2.4.1.115]
ko:K13080 C12RT1; flavanone 7-O-glucoside 2''-O-beta-L-rhamnosyltransferase [EC:2.4.1.236]
ko:K13311 desVI; dTDP-3-amino-3,4,6-trideoxy-alpha-D-glucopyranose N,N-dimethyltransferase [EC:2.1.1.234]
ko:K13921 pduQ; 1-propanol dehydrogenase
ko:K13937 H6PD; hexose-6-phosphate dehydrogenase [EC:1.1.1.47 3.1.1.31]
ko:K16266 UGT85K; cyanohydrin UDP-glucosyltransferase [EC:2.4.1.-]
ko:K16839 hpxO; FAD-dependent urate hydroxylase [EC:1.14.13.113]
ko:K16969 msmB; methanesulfonate monooxygenase subunit beta [EC:1.14.13.111]
ko:K17399 DNMT3B; DNA (cytosine-5)-methyltransferase 3B [EC:2.1.1.37]
ko:K18368 CSE; caffeoylshikimate esterase [EC:3.1.1.-]
ko:K19964 HIUH; hydroxyisourate hydrolase [EC:3.5.2.17]
ko:K21064 ycsE; 5-amino-6-(5-phospho-D-ribitylamino)uracil phosphatase [EC:3.1.3.104]
ko:K21202 2-ODD; (-)-deoxypodophyllotoxin synthase [EC:1.14.20.8]

ko:K22395 cinnamyl-alcohol dehydrogenase [EC:1.1.1.195]
ko:K22846 CS26; S-sulfo-L-cysteine synthase (O-acetyl-L-serine-dependent) [EC:2.5.1.144]
ko:K23094 ABC4; 2-carboxy-1,4-naphthoquinone phytyltransferase [EC:2.5.1.130]
ko:K26495 UGT91AP1; beta-1,3-glucosyltransferase [EC:2.4.1.-]

map01110 Biosynthesis of secondary metabolites (27)
ko:K00487 CYP73A; trans-cinnamate 4-monooxygenase [EC:1.14.14.91]
ko:K00827 AGXT2; alanine-glyoxylate transaminase / (R)-3-amino-2-methylpropionate-pyruvate transaminase [EC:2.6.1.44 2.6.1.40]
ko:K00901 dgkA; diacylglycerol kinase (ATP) [EC:2.7.1.107]
ko:K00939 adk; adenylate kinase [EC:2.7.4.3]
ko:K00944 AK3; nucleoside-triphosphate--adenylate kinase [EC:2.7.4.10]
ko:K01092 E3.1.3.25; myo-inositol-1(or 4)-monophosphatase [EC:3.1.3.25]
ko:K04123 KAO; ent-kaurenoic acid monooxygenase [EC:1.14.14.107]
ko:K06134 COQ7; 3-demethoxyubiquinol 3-hydroxylase [EC:1.14.99.60]
ko:K06900 capV; cGAMP-activated phospholipase [EC:3.1.1.32 3.1.1.-]
ko:K08249 E4.1.2.11; hydroxymandelonitrile lyase [EC:4.1.2.11]
ko:K12355 REF1; coniferyl-aldehyde dehydrogenase [EC:1.2.1.68]
ko:K12930 BZ1; anthocyanidin 3-O-glucosyltransferase [EC:2.4.1.115]
ko:K13071 PAO; pheophorbide a oxygenase [EC:1.14.15.17]
ko:K13311 desVI; dTDP-3-amino-3,4,6-trideoxy-alpha-D-glucopyranose N,N-dimethyltransferase [EC:2.1.1.234]
ko:K13492 ZOG1; zeatin O-glucosyltransferase [EC:2.4.1.203]
ko:K13496 UGT73C; UDP-glucosyltransferase 73C [EC:2.4.1.-]
ko:K13937 H6PD; hexose-6-phosphate dehydrogenase [EC:1.1.1.47 3.1.1.31]
ko:K15095 E1.1.1.208; (+)-neomenthol dehydrogenase [EC:1.1.1.208]
ko:K15404 CER1; aldehyde decarbonylase [EC:4.1.99.5]
ko:K16085 CYP99A2_3; 9beta-pimara-7,15-diene oxidase [EC:1.14.14.111]
ko:K16266 UGT85K; cyanohydrin UDP-glucosyltransferase [EC:2.4.1.-]
ko:K18368 CSE; caffeoylshikimate esterase [EC:3.1.1.-]
ko:K21064 ycsE; 5-amino-6-(5-phospho-D-ribitylamino)uracil phosphatase [EC:3.1.3.104]
ko:K21202 2-ODD; (-)-deoxypodophyllotoxin synthase [EC:1.14.20.8]
ko:K22395 cinnamyl-alcohol dehydrogenase [EC:1.1.1.195]
ko:K23094 ABC4; 2-carboxy-1,4-naphthoquinone phytyltransferase [EC:2.5.1.130]
ko:K26495 UGT91AP1; beta-1,3-glucosyltransferase [EC:2.4.1.-]

map04626 Plant-pathogen interaction (11)
ko:K13416 BAK1; brassinosteroid insensitive 1-associated receptor kinase 1 [EC:2.7.10.1 2.7.11.1]
ko:K13420 FLS2; LRR receptor-like serine/threonine-protein kinase FLS2 [EC:2.7.11.1]
ko:K13430 PBS1; serine/threonine-protein kinase PBS1 [EC:2.7.11.1]

ko:K13434 PTI6; pathogenesis-related genes transcriptional activator PTI6
ko:K13436 PTI1; pto-interacting protein 1 [EC:2.7.11.1]
ko:K13448 CML; calcium-binding protein CML
ko:K13453 PRF; disease resistance protein
ko:K13457 RPM1; disease resistance protein RPM1
ko:K13459 RPS2; disease resistance protein RPS2
ko:K13460 RPS5; disease resistance protein RPS5
ko:K18878 UPA20; BHLH transcription factor Upa20

map04075 Plant hormone signal transduction (9)
ko:K12126 PIF3; phytochrome-interacting factor 3
ko:K13415 BRI1; protein brassinosteroid insensitive 1 [EC:2.7.10.1 2.7.11.1]
ko:K13416 BAK1; brassinosteroid insensitive 1-associated receptor kinase 1 [EC:2.7.10.1 2.7.11.1]
ko:K13422 MYC2; transcription factor MYC2
ko:K13464 JAZ; jasmonate ZIM domain-containing protein
ko:K14486 ARF; auxin response factor
ko:K14494 DELLA; DELLA protein
ko:K14495 GID2; F-box protein GID2
ko:K14510 CTR1; serine/threonine-protein kinase CTR1 [EC:2.7.11.1]

map04120 Ubiquitin mediated proteolysis (6)
ko:K03175 TRAF6; TNF receptor-associated factor 6 [EC:2.3.2.27]
ko:K03361 CDC4; F-box and WD-40 domain protein CDC4
ko:K08770 UBC; ubiquitin C
ko:K10573 UBE2A; ubiquitin-conjugating enzyme E2 A [EC:2.3.2.23]
ko:K10594 HERC1; E3 ubiquitin-protein ligase HERC1 [EC:2.3.2.26]
ko:K10699 UBE1L2; ubiquitin-activating enzyme E1-like protein 2 [EC:6.2.1.45]

map04712 Circadian rhythm - plant (6)
ko:K12121 PHYB; phytochrome B
ko:K12126 PIF3; phytochrome-interacting factor 3
ko:K12130 PRR5; pseudo-response regulator 5
ko:K12135 CO; zinc finger protein CONSTANS
ko:K16221 TCP21; transcription factor TCP21 (protein CCA1 HIKING EXPEDITION)
ko:K16222 CDF1; Dof zinc finger protein DOF5.5

map04814 Motor proteins (5)

ko:K09290 TPM3; tropomyosin 3
ko:K10357 MYO5; myosin V

ko:K10395 KIF4; kinesin family member 4
ko:K10396 KIF5; kinesin family member 5
ko:K10412 DNAL4; dynein axonemal light chain 4

map00130 Ubiquinone and other terpenoid-quinone biosynthesis (3)
ko:K00487 CYP73A; trans-cinnamate 4-monooxygenase [EC:1.14.14.91]
ko:K06134 COQ7; 3-demethoxyubiquinol 3-hydroxylase [EC:1.14.99.60]
ko:K23094 ABC4; 2-carboxy-1,4-naphthoquinone phytyltransferase [EC:2.5.1.130]

map00380 Tryptophan metabolism (1)
ko:K23947 DAO; 2-oxoglutarate-dependent dioxygenase [EC:1.14.11.-]

**Stable 8. Pathways and links of DEGs identified between stem and root**

#### Listed links of stem_vs_root DEGs involved pathways

map01100 Metabolic pathways (18)
ko:K00333 nuoD; NADH-quinone oxidoreductase subunit D [EC:7.1.1.2]
ko:K00434 E1.11.1.11; L-ascorbate peroxidase [EC:1.11.1.11]
ko:K00827 AGXT2; alanine-glyoxylate transaminase / (R)-3-amino-2-methylpropionate-pyruvate transaminase [EC:2.6.1.44 2.6.1.40]
ko:K00876 udk; uridine kinase [EC:2.7.1.48]
ko:K01100 E3.1.3.37; sedoheptulose-bisphosphatase [EC:3.1.3.37]
ko:K01771 plc; 1-phosphatidylinositol phosphodiesterase [EC:4.6.1.13]
ko:K02641 petH; ferredoxin--NADP+ reductase [EC:1.18.1.2]
ko:K05714 mhpC; 2-hydroxy-6-oxonona-2,4-dienedioate hydrolase [EC:3.7.1.14]
ko:K06900 capV; cGAMP-activated phospholipase [EC:3.1.1.32 3.1.1.-]
ko:K08729 PTDSS1; phosphatidylserine synthase 1 [EC:2.7.8.-]
ko:K11419 SUV39H; [histone H3]-lysine9 N-trimethyltransferase SUV39H [EC:2.1.1.355]
ko:K12930 BZ1; anthocyanidin 3-O-glucosyltransferase [EC:2.4.1.115]
ko:K13311 desVI; dTDP-3-amino-3,4,6-trideoxy-alpha-D-glucopyranose N,N-dimethyltransferase [EC:2.1.1.234]
ko:K13937 H6PD; hexose-6-phosphate dehydrogenase [EC:1.1.1.47 3.1.1.31]
ko:K16839 hpxO; FAD-dependent urate hydroxylase [EC:1.14.13.113]
ko:K16969 msmB; methanesulfonate monooxygenase subunit beta [EC:1.14.13.111]
ko:K21064 ycsE; 5-amino-6-(5-phospho-D-ribitylamino)uracil phosphatase [EC:3.1.3.104]
ko:K23094 ABC4; 2-carboxy-1,4-naphthoquinone phytyltransferase [EC:2.5.1.130]

map01110 Biosynthesis of secondary metabolites (10)
ko:K00827 AGXT2; alanine-glyoxylate transaminase / (R)-3-amino-2-methylpropionate-pyruvate transaminase [EC:2.6.1.44 2.6.1.40]

ko:K06900 capV; cGAMP-activated phospholipase [EC:3.1.1.32 3.1.1.-]
ko:K12930 BZ1; anthocyanidin 3-O-glucosyltransferase [EC:2.4.1.115]
ko:K13071 PAO; pheophorbide a oxygenase [EC:1.14.15.17]
ko:K13311 desVI; dTDP-3-amino-3,4,6-trideoxy-alpha-D-glucopyranose N,N-dimethyltransferase [EC:2.1.1.234]
ko:K13937 H6PD; hexose-6-phosphate dehydrogenase [EC:1.1.1.47 3.1.1.31]
ko:K16085 CYP99A2_3; 9beta-pimara-7,15-diene oxidase [EC:1.14.14.111]
ko:K21026 AAE; acetylajmaline esterase [EC:3.1.1.80]
ko:K21064 ycsE; 5-amino-6-(5-phospho-D-ribitylamino)uracil phosphatase [EC:3.1.3.104]
ko:K23094 ABC4; 2-carboxy-1,4-naphthoquinone phytyltransferase [EC:2.5.1.130]

map04626 Plant-pathogen interaction (9)
ko:K13416 BAK1; brassinosteroid insensitive 1-associated receptor kinase 1 [EC:2.7.10.1 2.7.11.1]
ko:K13420 FLS2; LRR receptor-like serine/threonine-protein kinase FLS2 [EC:2.7.11.1]
ko:K13430 PBS1; serine/threonine-protein kinase PBS1 [EC:2.7.11.1]
ko:K13434 PTI6; pathogenesis-related genes transcriptional activator PTI6
ko:K13436 PTI1; pto-interacting protein 1 [EC:2.7.11.1]
ko:K13448 CML; calcium-binding protein CML
ko:K13453 PRF; disease resistance protein
ko:K13457 RPM1; disease resistance protein RPM1
ko:K18878 UPA20; BHLH transcription factor Upa20

map04075 Plant hormone signal transduction (6)
ko:K13415 BRI1; protein brassinosteroid insensitive 1 [EC:2.7.10.1 2.7.11.1]
ko:K13416 BAK1; brassinosteroid insensitive 1-associated receptor kinase 1 [EC:2.7.10.1 2.7.11.1]
ko:K13422 MYC2; transcription factor MYC2
ko:K13464 JAZ; jasmonate ZIM domain-containing protein
ko:K14491 ARR-B; two-component response regulator ARR-B family
ko:K14494 DELLA; DELLA protein

map04016 MAPK signaling pathway - plant (3)
ko:K13416 BAK1; brassinosteroid insensitive 1-associated receptor kinase 1 [EC:2.7.10.1 2.7.11.1]
ko:K13420 FLS2; LRR receptor-like serine/threonine-protein kinase FLS2 [EC:2.7.11.1]
ko:K13422 MYC2; transcription factor MYC2

map01120 Microbial metabolism in diverse environments (3)
ko:K01100 E3.1.3.37; sedoheptulose-bisphosphatase [EC:3.1.3.37]
ko:K05714 mhpC; 2-hydroxy-6-oxonona-2,4-dienedioate hydrolase [EC:3.7.1.14]

_ko:K16839 hpxO; FAD-dependent urate hydroxylase [EC:1.14.13.113]

map04712 Circadian rhythm - plant (2)
_ko:K12135 CO; zinc finger protein CONSTANS
_ko:K16221 TCP21; transcription factor TCP21 (protein CCA1 HIKING EXPEDITION)

map00130 Ubiquinone and other terpenoid-quinone biosynthesis (1)
_ko:K23094 ABC4; 2-carboxy-1,4-naphthoquinone phytyltransferase [EC:2.5.1.130]

#### 2. Supplementary Figures

##########################################################################

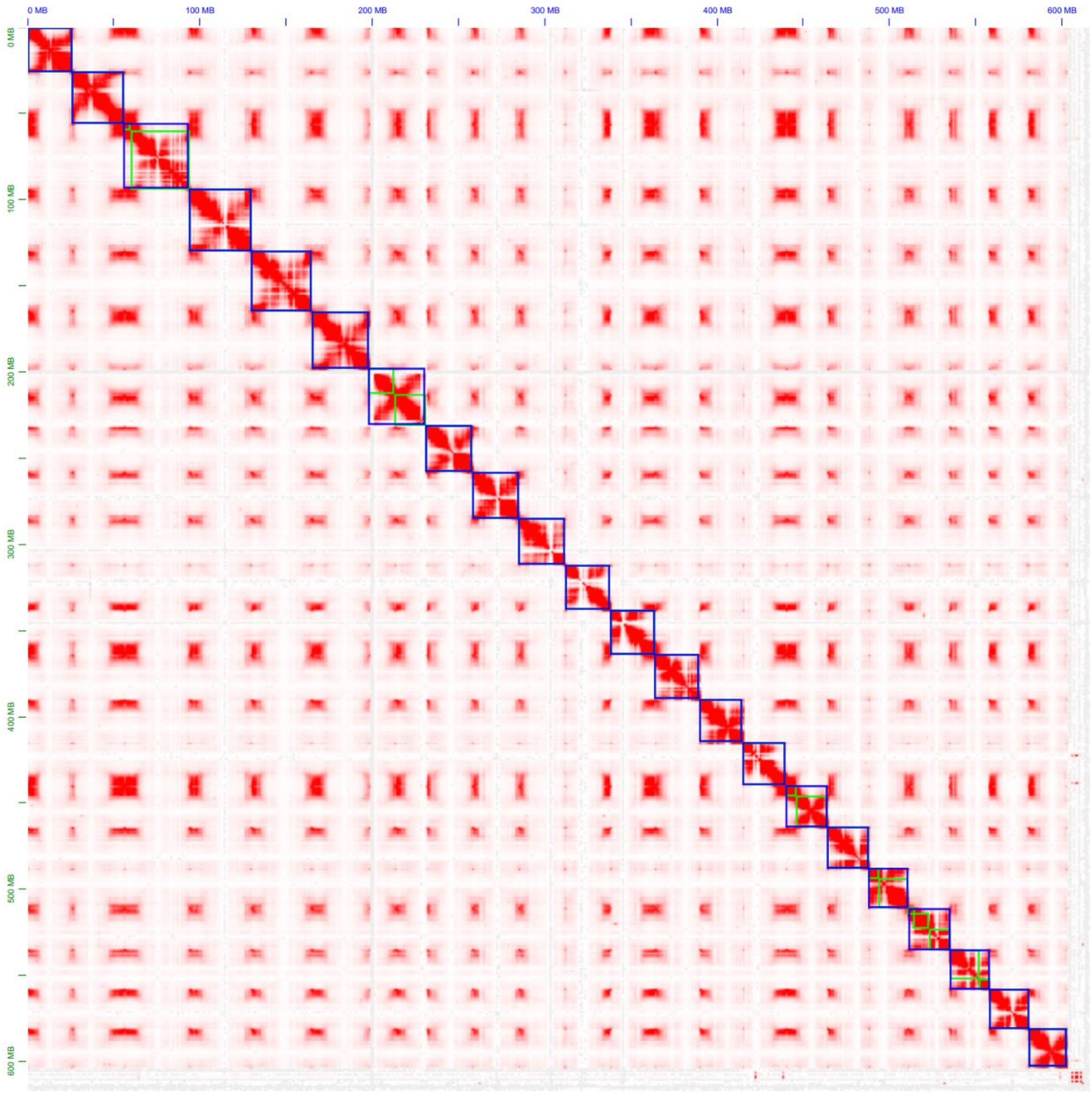

**Sfigure 1. HiC-map of hap1 showing chromosomal scaffolding**

Each blue box represents a chromosome, each green box represents the contigs.

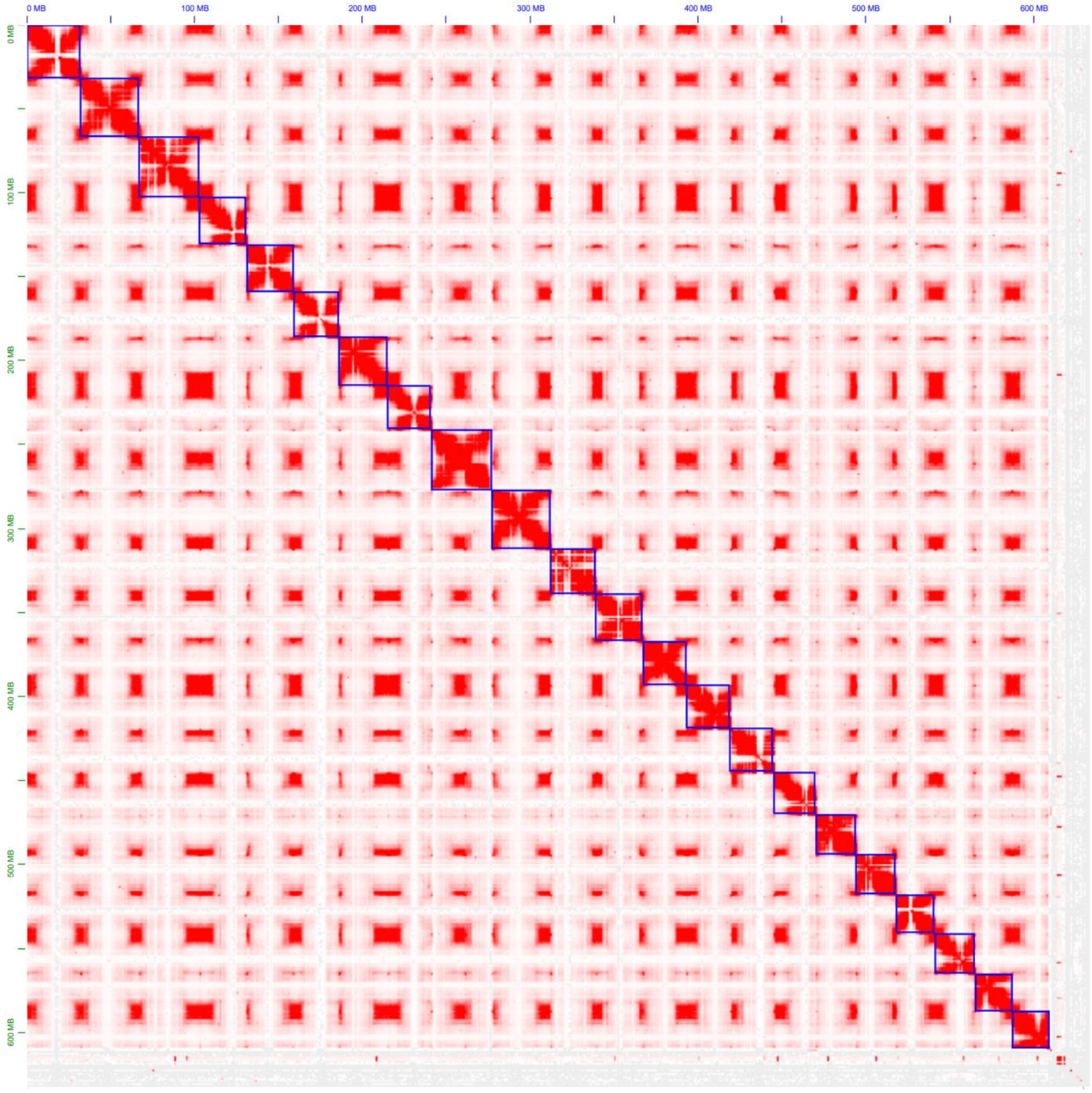

**Sfigure 2. HiC-map of hap2 showing chromosomal scaffolding**

Each blue box represents a chromosome, each green box represents the contigs.

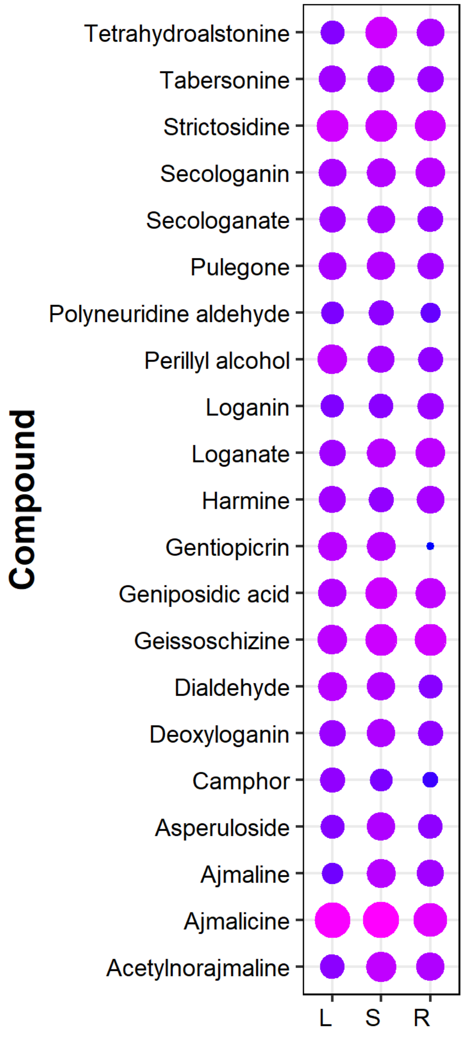

**Sfigure 3. Alkaloids abundance in tissues in the map 00902 and 00901**

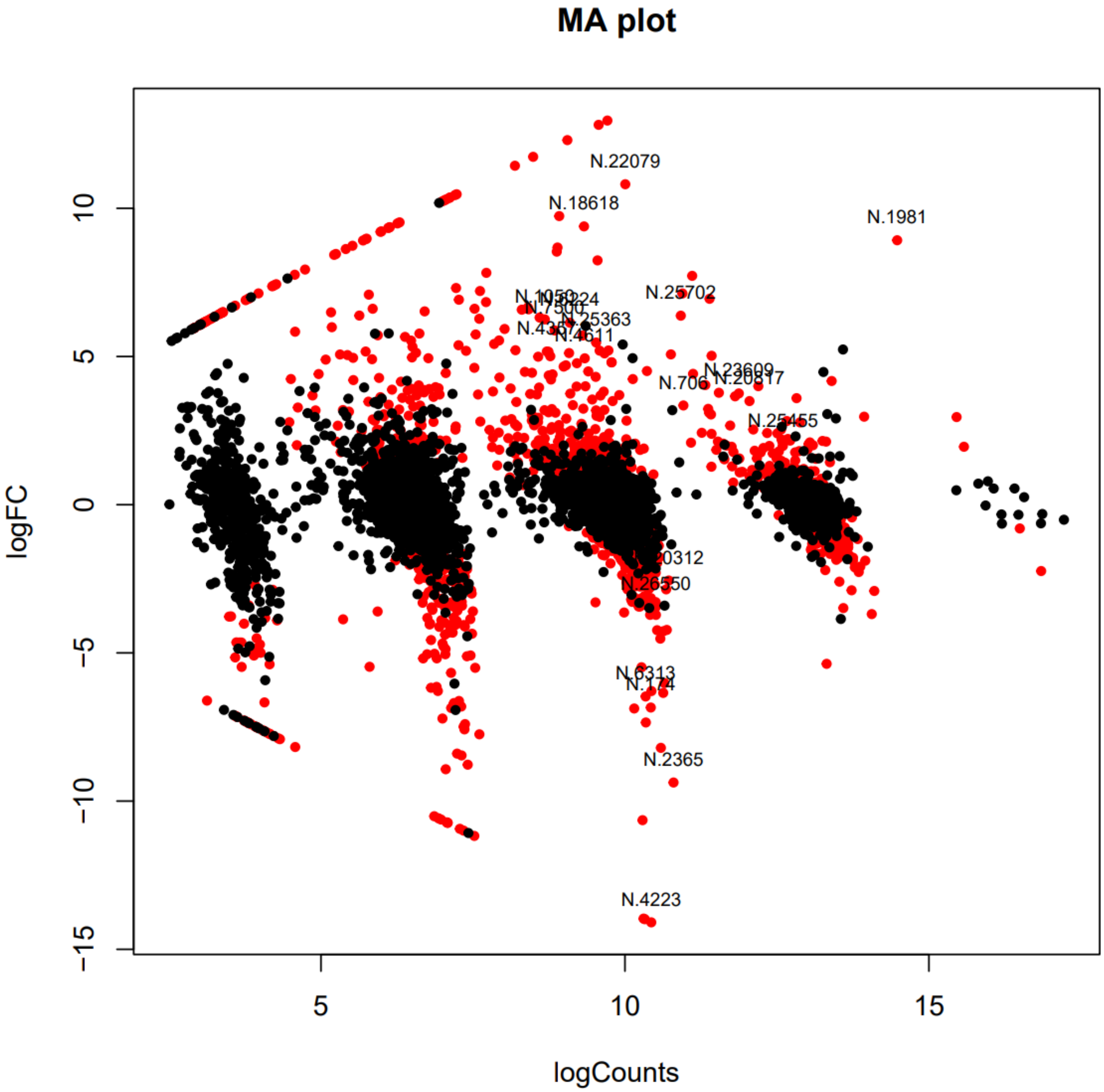

**Sfigure 4. Sfigure 2. MA plot showing DEGs between stem and root (control)**

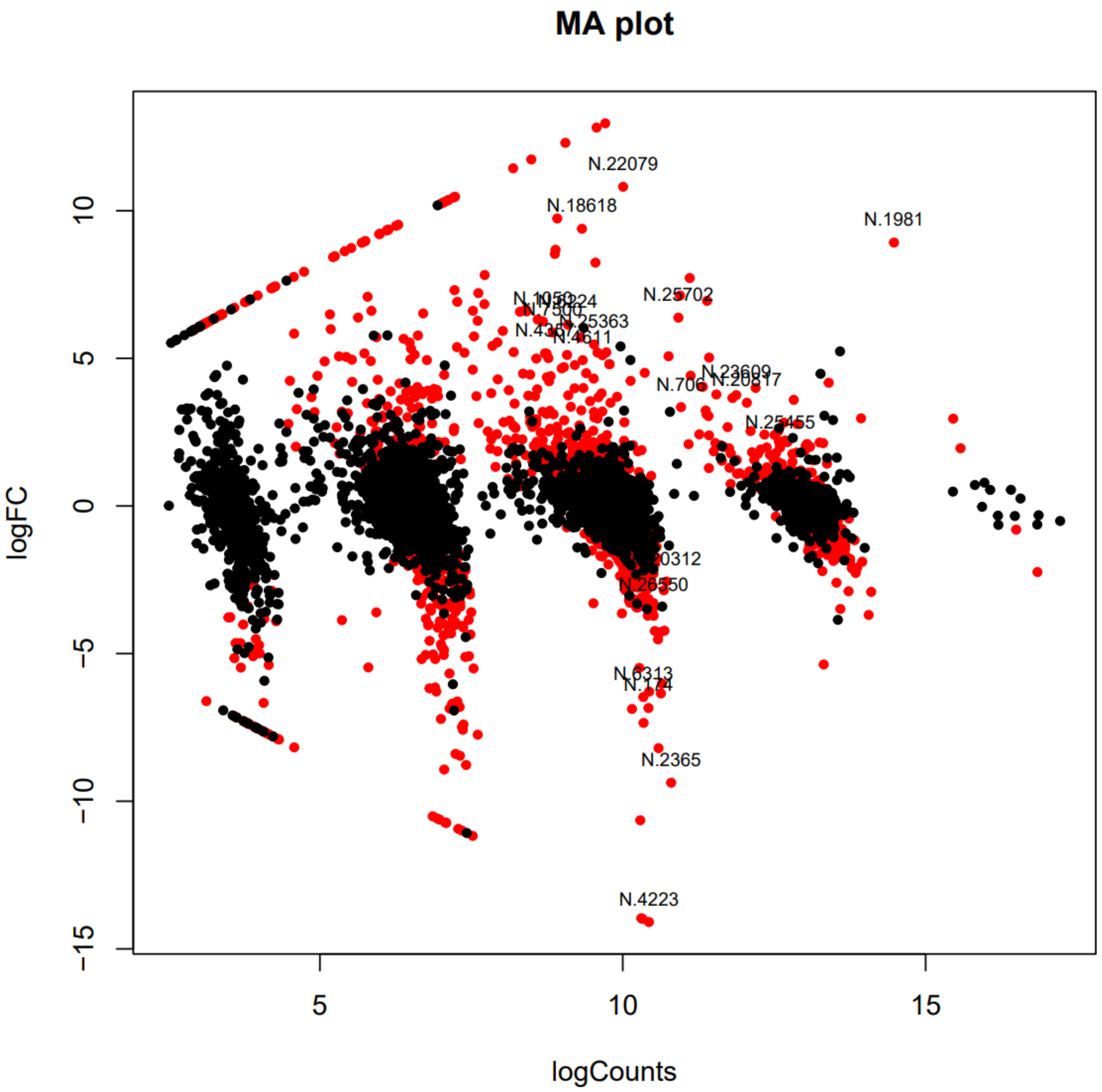

**Sfigure 5. MA plot showing DEGs between stem and leaf (control)**

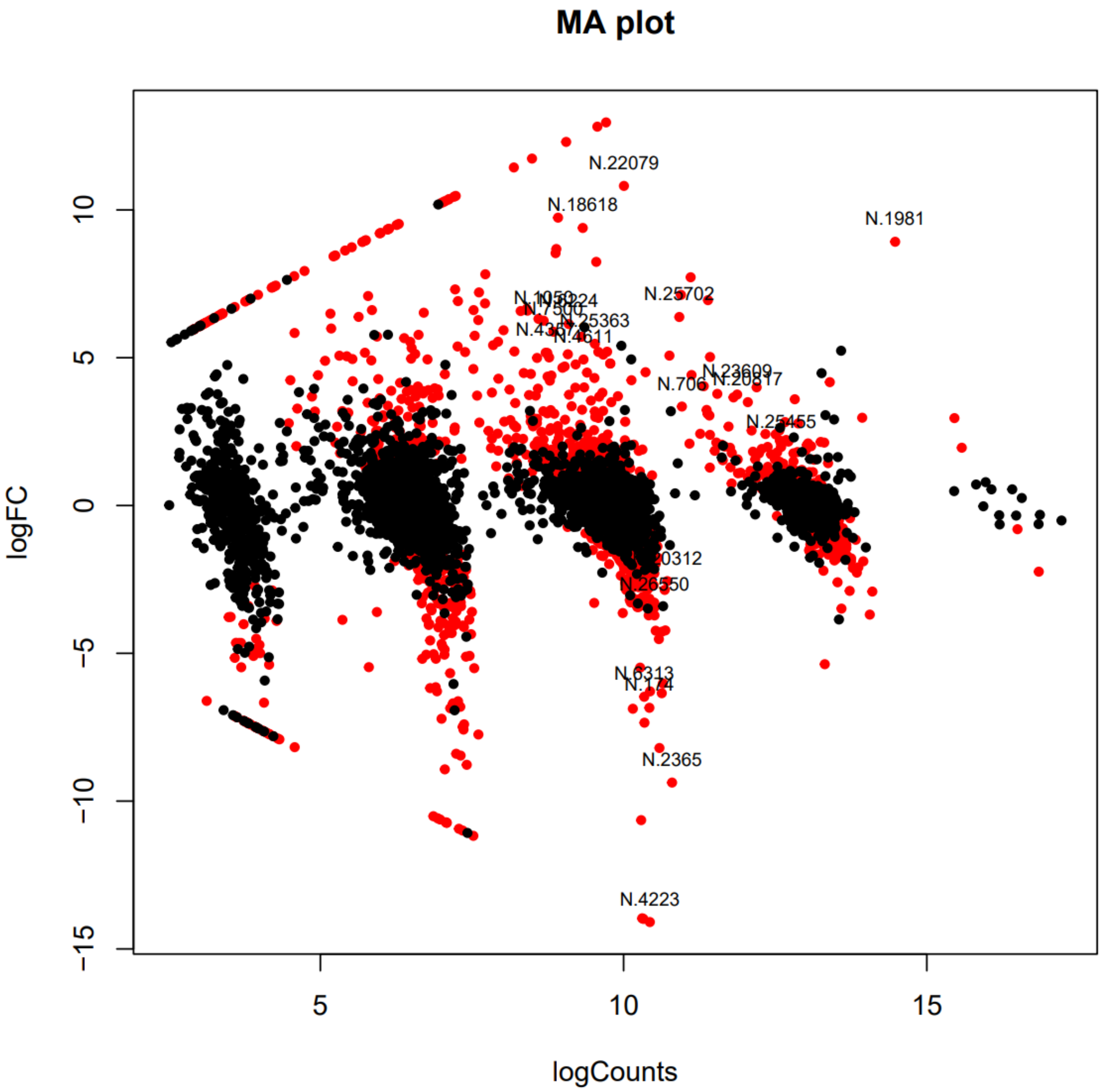

**Sfigure 6. MA plot showing DEGs between leaf and root (control)**
